# Genetic interactions of histone acetyl-transferase enzymes encoding genes *Gcn5* and *Mof* with *hsrω* lncRNA gene

**DOI:** 10.1101/509737

**Authors:** Deo Prakash Chaturvedi

## Abstract

The hsrω lncRNAs are known to interact with the Iswi chromatin remodeler while Iswi is known to interact with Gcn5, a general histone acetyl transferase, and Mof, a male-specific HAT essential for H4K16 acetylation and consequent hyperactivity of the single X-chromosome in male *Drosophila*. We show here that *hsrω* genetically interacts with Gcn5 as well as Mof, but unlike the suppression of phenotypes due to down-regulation or absence of Iswi, those following down-regulation of Gcn5 or Mof are suppressed by over-expression of *hsrω*. General lethality caused by *Act-GAL4* driven global expression of *Gcn5-RNAi* and the male-specific lethality following *Mof-RNAi* transgene expression were partially suppressed by over-expression of *hsrω*, but not by down regulation through *hsrω-RNAi*. Likewise, eye phenotypes following *ey-GAL4* driven down-regulation of Gcn5 or Mof were also partially suppressed by over-expression of *hsrω. Act-GAL4* driven global over-expression of *hsrω* along with *Gcn5-RNAi* transgene substantially restored levels of Gcn5 RNA as well as protein that were reduced by Gcn5-RNAi. *Mof-RNAi* transgene expression reduced Megator and Msl-2 levels and their nuclear distribution patterns; over-expression of *hsrω* along with *Mof-RNAi* substantially restored Megator levels and its distribution at the nuclear rim and in nucleoplasmic speckles and at the same time restored the male X-chromosome specific localization of Msl-2. Earlier reported antagonistic interactions of Mof with Iswi and interaction of hsrω transcripts with Megator appear to underlie the suppression of Gcn5 and Mof phenotypes by over-expression of the lncRNAs. Present results add the dosage compensation pathway to the list of diverse pathways in which the multiple lncRNAs produced by the *hsrω* are known to have important roles.

## Introduction

The cell type specific expression of genes, in spite of their apparently common genome, involves multiple regulatory mechanisms operating at transcriptional, post-transcriptional, translational and post-translational levels. Post-translational modifications of histones are now known to be essential for tissue specific gene expression patterns in eukaryotes (Kuo and Allis 1998; Kimura *et al*. 2005; Shahbazian and Grunstein 2007; Venkatesh and Workman 2015). Among the several modifications, histone acetylation plays a crucial role in activation of genes. The p55 of *Tetrahymena* was the first known histone acetyl transferase (HAT) (Brownell and Allis 1995) and was later found to be a homologue of the eukaryotic <u>G</u>eneral <u>C</u>ontrol <u>N</u>on-derepressible 5 or Gcn5 (Smith *et al*. 1998). Higher eukaryotes have three families of HATs, viz., GNAT, MYST and p300/CBP (Marmorstein and Trievel 2009). Gcn5 is a member of the GNAT family (Kuo and Allis 1998) and like other HATs, it also works as the acetyl transferase subunit of multicomponent ATAC (<u>A</u>da <u>T</u>wo <u>A</u> <u>C</u>ontaining) and SAGA (<u>S</u>pt-<u>A</u>da-<u>Gcn5-A</u>cetyl transferase) complexes (Guelman *et al*. 2006; Suganuma *et al*. 2008). Gcn5 is remarkably conserved through the metazoan phyla, being represented by a single gene in most metazoans, although vertebrates have two paralogous genes (Nagy and Tora 2007).

Male absent on first (Mof) is also a HAT, representing the MYST family in *Drosophila*. Mof, as a well known member of the *Drosophila* dosage compensation complex (DCC) (Hilfiker *et al*. 1997; Kind *et al*. 2008) acetylates H4K16 on male X chromosome to make it hyperactive (Conrad and Akhtar 2012).

The *hsrω* is a developmentally active and stress responsive essential gene which produces multiple long non-coding RNAs (lncRNA); its nuclear transcripts that contain the 280bp tandem repeats are involved in organization of the nucleoplasmic omega speckles (Prasanth *et al*. 2000, Lakhotia 2011, 2012, 2016). It has been shown that organization of omega speckles requires the ATP-dependent chromatin remodeling protein, Iswi (Onorati *et al*. 2011; Singh and Lakhotia 2015). Iswi is well known for the maintenance of chromatin architecture as its absence results in chromosome decondensation and late larval lethality (Corona and Tamkun 2004). Expression of *hsrω-RNAi* transgene, which results in down-regulation of the 280bp repeat containing nuclear transcripts of *hsrω* (Mallik and Lakhotia 2009), in Iswi null background was shown to partially restore chromatin architecture and suppress the larval lethality (Onorati *et al*. 2011). Iswi has also been shown to interact with DCC components like Mle and Mof (Corona *et al*. 2002) and with Gcn5 (Ferreira *et al*. 2007). The chromatin organization of the hyperactive male X chromosome in *Drosophila* is similarly affected by loss of Iswi or Gcn5 (Corona *et al*. 2002; Carre *et al*. 2008; Onorati *et al*. 2011). Interestingly, expression of *hsrω-RNAi* transgene in Iswi null background was found to restore the male X chromosome morphology (Onorati *et al*. 2011). In view of such interactions between *hsrω* and Iswi on one hand, and between Iswi and Gcn5 and Mof on the other, we have examined if HATs like Gcn5and Mof also interact with *hsrω*.

We show that the ATP-independent Gcn5 and Mof HATs genetically interact with hsrω transcripts since lethality and some other phenotypes caused by the down-regulation of Gcn5 or Mof are partially suppressed by over-expression of *hsrω*.

## Materials and methods

### Fly strains

All flies were maintained on standard cornmeal-agar food medium at 24±1°C. Oregon R+ strain was used as wild type (WT). For down-regulation of Gcn5 and Mof, *Gcn5-RNAi* (Bloomington Stock Center # 33981) and *Mof-RNAi* (Bloomington Stock Center # 31401) transgenic stocks, respectively, were used. The *w; +/+; UAS-hsrω-RNAi^3^/UAS-hsrω-RNAi^3^* transgenic line expressing both strands of the 280 bp repeat unit of the *hsrω* gene when driven by a *GAL4* driver was used for down-regulation of the repeat containing nuclear transcripts of *hsrω* gene (Mallik and Lakhotia 2009). This transgene is inserted on chromosome 3 and is referred to here as *hsrω-RNAi*. For over-expression of the *hsrω*, we used two *EP* alleles (Brand and Perrimon 1993) of *hsrω*, viz. *w; EP3037/EP3037* and *w; EP93D/EP93D* (Mallik and Lakhotia 2009). As described by Mallik and Lakhotia (2009), these lines carry an EP element (Brand and Perrimon 1993) at - 144 and −130 position, respectively, from the *hsrω* gene’s major transcription start site (www.flybase.org), resulting in over-expression of hsrω transcripts when driven by a *GAL4* driver. We used either the globally expressed *Actin5C-GAL4* (Ekengren *et al*. 2001) or the larval salivary gland (SG) and eye disc specific *eyeless-GAL4* (Halder *et al*. 1995) to drive expression of the target *RNAi transgene* or the *EP* alleles. These drivers are referred to as *Act-GAL4* and *ey-GAL4*, respectively in the text as well as in images. Appropriate crosses were made to obtain progenies of the desired genotypes.

### SDS-PAGE and western blotting

Wandering late 3^rd^ instar larvae (approx. 115 hrs after egg laying) of the desired genotype were dissected in Poels’ salt solution (PSS, Lakhotia and Tapadia 1998) and the internal tissues were immediately transferred to boiling sodium dodecyl sulfate (SDS) sample buffer (Laemmli 1970) for 10 min. Following lysis, the larval proteins were separated by SDS-polyacrylamide gel electrophoresis (SDS-PAGE) followed by western blotting as described earlier (Prasanth *et al*. 2000). The primary antibodies used were rabbit anti-H3K9Ac (9671 at 1:1000 dilution, Cell Signaling Technology, USA), and mouse anti-β-tubulin (E7 at 1:100 dilution, Sigma-Aldrich, India). Corresponding secondary antibodies were HRP-tagged anti-rabbit, and anti-mouse (1:1500 dilution, Bangalore Genei, India), respectively. The signal was detected using the Supersignal West Pico Chemiluminescent Substrate kit (Pierce, USA). Each western blotting was repeated at least twice. For reprobing a blot with another antibody after detection of the first antibody binding was completed, the blot membrane was kept in stripping buffer (100 mM 2-mercaptoethanol, 2 % SDS, 62.5 mM TrisHCl pH 6.8) at 50 °C for 30 min on a shaker bath followed by processing for detection with the desired second antibody.

### Tissue immunofluorescence

Salivary glands (SG) or Malpighian tubules (MT) from wandering late 3^rd^ instar larvae (approx 115 hrs after egg laying) of the desired genotypes were dissected out in PSS and processed for immunostaining as described earlier (Prasanth *et al*. 2000). Primary antibodies used were rabbit anti-Msl-2 (1:100 dilution, Strukov *et al*. 2011; Graindorge *et al*. 2013), rabbit anti-H3K9Ac (1:100 dilution, Cell Signaling Technology, USA) and mouse anti-Megator BX34 (1:20 dilution, Zimowska and Paddy 2002). Cy3 (1:200 dilution, Sigma-Aldrich, India) or Alexa Flour (1:200 dilution, Molecular Probe) conjugated anti-rabbit or anti-mouse antibodies were used as secondary antibodies as required. The immunostained tissues were counterstained with DAPI, mounted in DABCO and examined under laser scanning confocal microscope, Zeiss LSM 510 Meta, using appropriate filters/dichroics required for the given fluorochrome. Each immunostaining was carried out at least twice. All images were assembled using Adobe Photoshop 7.0.

### RNA isolation and RT-PCR

Total RNA was isolated from late 3^rd^ instar larvae of the desired genotypes using the TRI reagent as per the manufacturer’s (Sigma-Aldrich, India) instructions. RNA pellets were resuspended in nuclease-free water. The cDNA was synthesized as described earlier (Lakhotia *et al*. 2012), and semi-quantitative RT-PCR was carried out for the desired transcripts. G3PDH cDNA, used as loading control, was PCR-amplified with 5’-CCACTGCCGAGGAGGTCAACTA-3 as the forward and 5’-GCTCAGGGTGATTGCGTATGCA-3’ as the reverse primers. The Gcn5 cDNA was amplified using 5’-CCAGTTTATGCGGGCTACAT-3’ as forward and 5’-CCCTCCTTGAAGCAAGTCAA-3’ as reverse primers. The thermal cycling parameters included an initial denaturation at 94° C for 5 min followed by 25 cycles of 30 s at 94° C, 30 s at 56° C, and 30 s at 72° C. Final extension was at 72° C for 10 min. The PCR products were electrophoresed on 2.0 % agarose gel with a 100-bp DNA ladder marker (Bangalore Genei, India). Each RT-PCR was carried out with three independently prepared RNA samples.

### Nail polish imprints

The nail polish imprints of eyes of adult flies of desired genotypes were prepared as described (Arya and Lakhotia 2006) and were examined using DIC optics on a Nikon Ellipse 800 microscope.

### Muscle preparation

For analyzing thoracic dorsal longitudinal muscles (DLMs), 3-4 day old flies of the desired genotypes were frozen in liquid nitrogen and their thoraces were cut sagitally with a sharp blade and dehydrated through series of 50%, 70%, 90% and absolute ethanol, keeping for 10 min in each. After the dehydration, they were cleared overnight in methyl salicylate, mounted in Canada balsam and observed in plane polarized light using a Nikon E800 microscope (Singh and Roy 2013).

## Results

### Over-expression of hsrω partially rescued pupal lethality and dorsal closure defect in pharates seen following ubiquitous down-regulation of Gcn5

Down-regulation of Gcn5 was achieved through a homozygous viable RNAi line (Bloomington *Drosophila* Stock Center stock # 33981), with the transgene inserted on chromosome 3. For down-regulation of *hsrω*, we used the earlier characterized *hsrω-RNAi^3^* transgene while for overexpression, we used the *EP3037* or *EP93D* alleles of *hsrω* (Mallik and Lakhotia 2009). To examine the developmental consequences of *Act-GAL4* driven ubiquitous expression of the *UAS-Gcn5-RNAi* transgene, eggs from different genotypes (Fig. 1a) were collected at two hour intervals and grown on standard food at 24°C. Data on the proportion of eggs that eclosed as adult flies are presented in Fig. 1a. In agreement with earlier report (Carre *et al*. 2005) that absence of Gcn5 results in lethality at pharate stage and defects in cuticle development during metamorphosis, present results (Fig. 1a) also showed that ubiquitous down-regulation of Gcn5 resulted in complete lethality between 3^rd^ instar larval and pharate stages. Co-expression of *hsrω-RNAi* transgene (Mallik and Lakhotia 2009) did not affect the pattern or extent of lethality following down-regulation of Gcn5. However, co-expression of the over-expressing *EP3037* or *EP93D* allele of *hsrω* gene (Mallik and Lakhotia 2009) with *Gcn5-RNAi* transgene resulted in a partial rescue of the lethality. *Act-GAL4* driven co-expression of *Gcn5-RNAi* transgene with the *EP93D* allele resulted (Fig. 1a) in eclosion of ~3.5% and ~5.5%, respectively, of the male and female pupae as adults (genotypes 5 and 6 in Fig. 1a). Similar results were obtained when *EP3037*, the other over-expression allele of *hsrω*, was co-expressed with *Gcn5-RNAi* (data not presented).

**Fig. 1.**
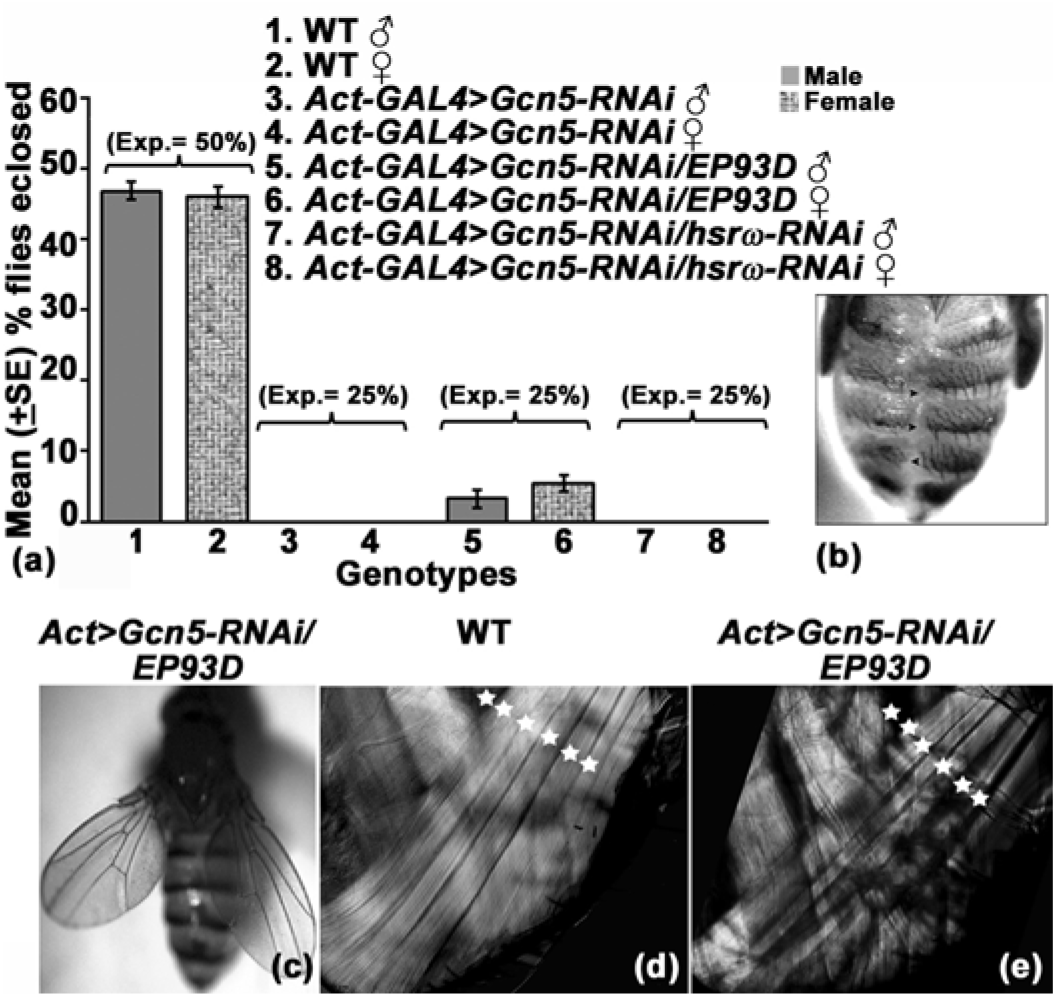
Lethality and dorsal closure defect during pupal metamorphosis following down regulation of Gcn5 are partially suppressed by over-expression of *hsrω*. **(a).** Graphical representation of mean % eclosion as adult flies (Y axis) of WT (N= 572, bars 1 and 2), *Act-GAL4>Gcn5-RNAi* (N= 564, bars 3 and 4), *Act-GAL4>Gcn5-RNAi/EP93D* (N= 550, bars 5 and 6) and *Act-GAL4>Gcn5-RNAi/hsrω-RNAi* (N= 544, bars 7 and 8). Expected frequencies (Exp) of adult male and female flies for different genotypes are noted above the histogram bars in parentheses. **(b).** Dorsal closure defect (arrowheads) seen in abdominal region of *Act-GAL4>Gcn5-RNAi* dead pharates. **(c).** *Act-GAL4>Gcn5-RNAi/EP93D* fly with normal dorsal closure but out-stretched wings. **(d)** and **(e).** Polarizing microscopic images of dorsal-longitudinal muscle fiber bundles (DLMs, white asterisks) in wild type and *Act-GAL4>Gcn5-RNAi/EP93D* flies, respectively.

About 90% of the dead pharates in the above crosses displayed dorsal closure defect in the middorsal region of abdomen (Fig. 1b). Frequency of dorsal closure defect in the *Gcn5-RNAi* expressing dead pharates (N = 143 for *Act-GAL4>Gcn5-RNAi* pharates, 172 for *Act-GAL4>Gcn5-RNAi/EP93D* pharates and 135 for *Act-GAL4>Gcn5-RNAi/hsrω-RNAi* pharates) remained about 90% irrespective of the levels of hsrω transcripts. Interestingly, however, none of the few emerging *Act-GAL4>Gcn5-RNAi/EP93D* flies showed the dorsal closure defect (Fig. 1c) although they were weak with stretched out wings (Fig. 1c) and unable to fly. They died within 2 weeks. Polarizing microscopic examination of the thoracic dorsal longitudinal muscle (DLM) fiber bundles in seven of these surviving flies revealed that six DLM bundles were present in each of the *Act-GAL4>Gcn5-RNAi/EP93D* thoraces like in wild type flies (Fig. 1d), they were improperly organized with greater inter-bundle spaces (Fig. 1e). Since no flies emerged when *Gcn5-RNAi* was expressed alone or in combination with *hsrω-RNAi* under the *Act-GAL4* driver, arrangement of DLMs in adults of these genotypes could not be examined. The muscle phenotype appears to be related to the reported role of Gcn5 mediated acetylation in muscle development (Sartorelli *et al*. 1999).

### Eye defects following *ey-GAL4* driven expression of *Gcn5-RNAi* were partially rescued by over-expression of hsrω

The *ey-GAL4* driven expression of *Gcn5-RNAi* transgene did not cause any lethality and all the pupae eclosed. However, their eyes, in all the flies, were smaller with ommatidial arrays being disordered (Fig. 2). Co-expression of *EP93D* allele (Fig. 2c, g), but not of the *hsrω-RNAi* transgene (Fig. 2d, h) substantially suppressed the small eye and disordered ommatidial arrangement in all the flies. Heads of *ey-GAL4>Gcn5-RNAi* and *ey-GAL4>Gcn5-RNAi/hsrω-RNAi* flies were also found to be smaller in comparison to wild type or *ey-GAL4>Gcn5-RNAi/EP93D* flies (not shown).

**Fig. 2.**
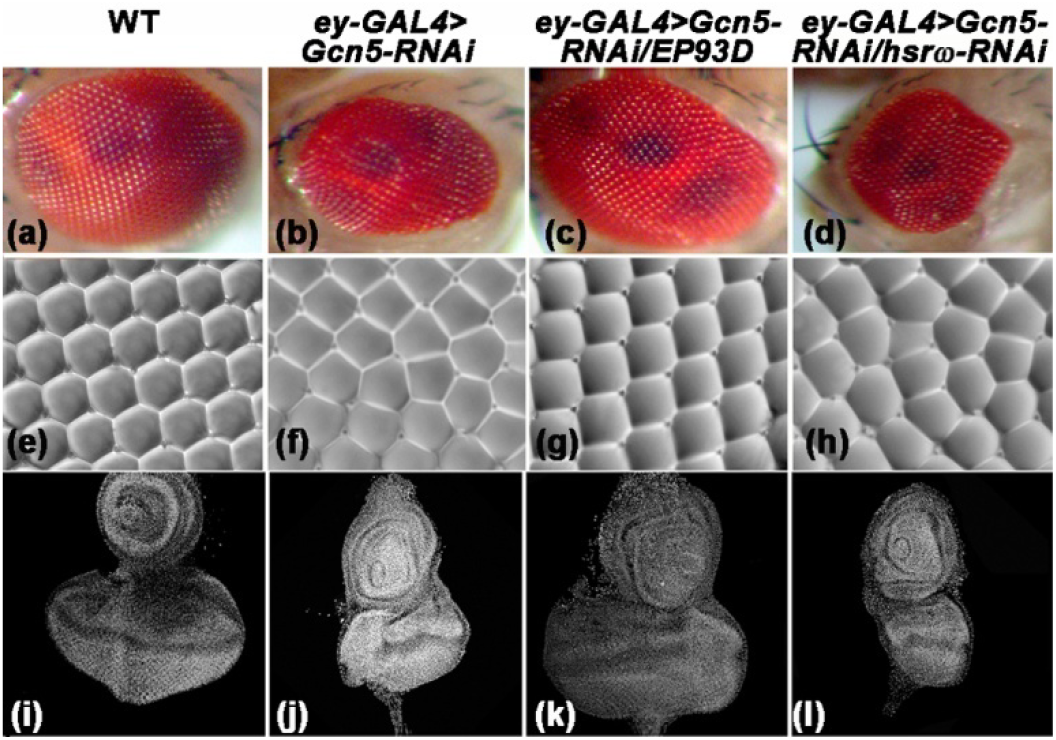
Down-regulation of Gcn5 with *ey-GAL4* driver affects eye development as seen in adult eye morphology (a-d), nail-polish imprints (e-h) and larval eye-antennal imaginal discs (i-l) of WT (a, e, i), *ey-GAL4>Gcn5-RNAi* (b, f, j), *ey-GAL4>Gcn5-RNAi/EP93D* (c, g, k) and *ey-GAL4>Gcn5-RNAi/hsrω-RNAi* (d, h, l).

In order to see if the eye-antennal imaginal discs were also affected by *Gcn5-RNAi* expression, eye discs from WT, *ey-GAL4>Gcn5-RNAi, ey-GAL4>Gcn5-RNAi/EP93D* and *ey-GAL4>Gcn5-RNAi/hsrω-RNAi* late 3^rd^ instar larvae were also examined after DAPI-staining. As shown in Fig. 2i-l, compared to the imaginal discs in wild type (Fig. 2i) larvae, those in any individuals in *ey-GAL4>Gcn5-RNAi* (Fig. 2j) and *ey-GAL4>Gcn5-RNAi/hsrω-RNAi* (Fig. 2l) larvae were much smaller. Interestingly, eye discs in all the *ey-GAL4>Gcn5-RNAi/EP93D* (Fig. 2k) larvae were nearly similar in size to those in wild type.

### Co-expression of EP93D transgene with *Gcn5-RNAi* restored Gcn5 transcripts to normaJ levels

Semi-quantitative RT-PCR data confirmed that, compared to WT, the Gcn5 transcript level was indeed reduced following *Act-GAL4* driven expression of *Gcn5-RNAi* transgene (Fig. 3c lane 2). However, following co-expression of the *EP93D* allele of *hsrω* (Fig. 3c lane 3), but not of the *hsrω-RNAi* transgene (Fig. 3c lane 4), the Gcn5 transcript levels were restored to the WT level. We also checked if ubiquitous expression of *EP93D* allele of *hsrω* or of the *hsrω-RNAi* transgene in *Gcn5^+^* background affected expression of Gcn5 (Fig. 3a). RT-PCR results showed that expression of Gcn5 was not affected by up- (Fig. 3a lane 2) or down-regulation (Fig. 3a lane 3) of hsrω transcripts in *Gcn5^+^* background.

**Fig. 3.**
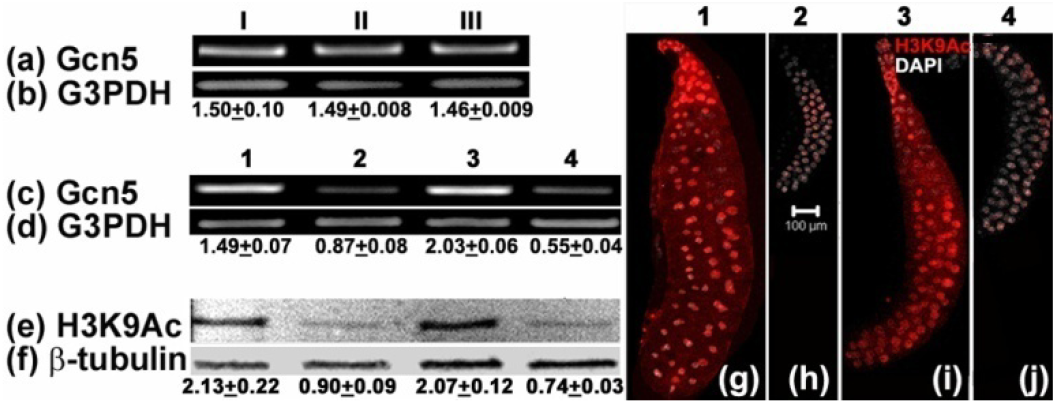
Images show ethedium bromide stained Gcn5 (a and c) and G3PDH (b and d) amplicons following semi-quantitative RT-PCR with total larval RNA. I, II and III in rows (a) and (b) represents genotypes *Act-GAL4, Act-GAL4>EP93D* and *Act-GAL4>hsrω-RNAi* respectively. 1, 2, 3 and 4 in images c-j represents genotypes WT, *Act-GAL4>Gcn5-RNAi, Act-GAL4>GCN5-RNAi/EP93D* and *Act-GAL4>GCN5-RNAi/hsrω-RNAi* respectively. Western blot showing levels of H3K9 acetylation in total proteins (e) from larvae of different genotypes; shown by 1-4 on top of each column, β-tubulin (f) was used as internal control. Values of mean relative fluorescence intensities (±S.E.) of Gcn5 and G3PDH bands (a-d) and H3K9Ac and β-tubulin bands (e and f) are given below the lanes b, d (N=3) and f (N=2). **(g-j)** represent confocal images showing nuclear distribution of K9 acetylated H3 (red) in whole salivary gland from larvae of different genotypes, shown by 1-4 on top of each column. Tissue was counterstained with DAPI (white) to localize nuclei. Scale bar in (h) represents 100μm and applies to g-j.

### Reduced H3K9 acetylation following down-regulation of Gcn5 is partially restored by coexpression of EP93D

Distribution and levels of H3K9 acetylated histones were examined in SG (Fig. 3g-j) from larvae of different genotypes. Immunostaining showed that compared to wild type cells (Fig. 3g), those expressing *Act-GAL4>Gcn5-RNAi* showed greatly reduced intensity of H3K9Ac (Fig. 3h). Acetylation of H3K9 was restored significantly following co-expression of *EP93D* allele of *hsrω* (Fig. 3i). However, the H3K9Ac levels in cells co-expressing *Gcn5-RNAi* and *hsrω-RNAi* transgenes (Fig. 3j) were as low as in those expressing only *Gcn5-RNAi*. Sizes of SG were also reduced following down-regulation of Gcn5 (Fig. 3h). Co-expression of *EP93D* allele of *hsrω* (Fig. 3i) but not *hsrω-RNAi* transgene (Fig. 3j) resulted in significant restoration of SG size.

In agreement with the above result, western blotting with anti-acetylated H3K9 showed that compared with levels in WT (Fig. 3e lane 1), down-regulation of Gcn5 reduced acetylation of H3K9 (Fig. 3e lane 2). Co-expression of *EP93D* allele of *hsrω* (Fig. 3e lane 3) but not the *hsrω-RNAi* transgene (Fig. 3e lane 4) with *Gcn5-RNAi* significantly restored H3K9 acetylation.

### Over-expression of hsrω transcripts rescued male lethality caused by global down-regulation of Mof

Down-regulation of Mof was achieved through expression of a homozygous viable *Mof-RNAi* transgene (stock # 31401 from the Bloomington *Drosophila* Stock Center) inserted on chromosome 3. To check the effect of global down-regulation of Mof on viability, the *UAS-Mof-RNAi* transgene was expressed ubiquitously using the *Act-GAL4* driver and data on survival to different developmental stages were collected. As reported earlier (Hilfiker *et al*. 1997), down-regulation of Mof resulted in complete lethality of males between 3^rd^ instar larval to pharate stage (Fig. 4a). Male lethality caused by the down-regulation of Mof was partially rescued by coexpression of *EP3037* allele of *hsrω* (Fig. 4a, genotype 5) but not by co-expression of *hsrω-RNAi* transgene (Fig. 4a, genotype 7), since about 2% male pupae (against the 12.5% expected) eclosed in the former case. Similar results were obtained when the *EP93D* over-expressing allele of *hsrω* was co-expressed with *Mof-RNAi* transgene (data not presented). Since, no lethality was seen in females (Fig. 4a), all further experiments were carried out only with males.

**Fig. 4.**
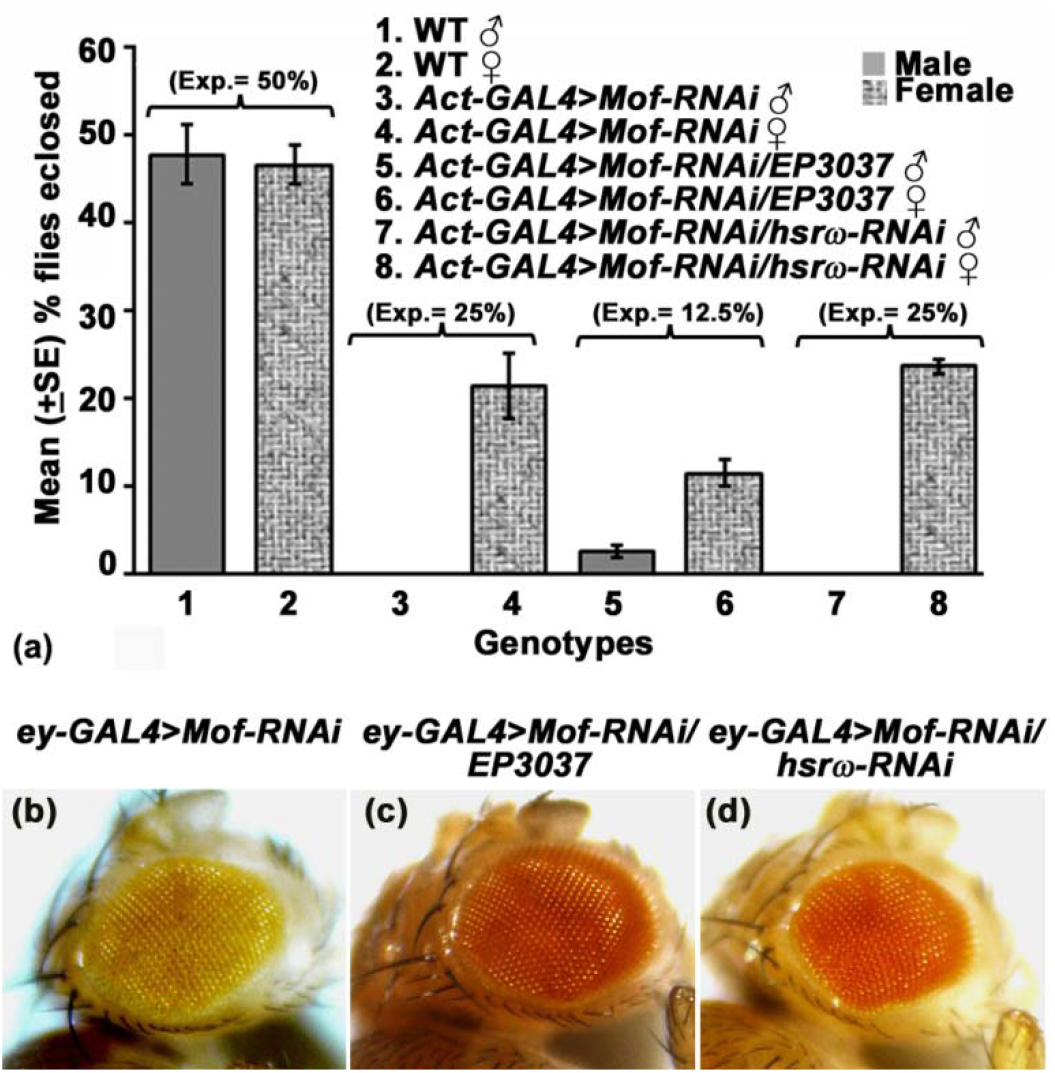
**(a)** Over-expression of *hsrω* partially suppresses male lethality and eye phenotypes resulting from global or eye-specific expression, respectively, of *Mof-RNAi* transgene. Graphical representation of mean % (±S.E.) eclosion of adult male and female flies (Y axis) of WT (N= 283, bars1 and 2), *Act-GAL4>Mof-RNAi* (N= 280, bars 3 and 4), *Act-GAL4>Mof-RNAi/EP3037* (N= 277, bars 5 and 6) and *Act-GAL4>Mof-RNAi/hsrω-RNAi* (N= 282, bars 7 and 8). Expected eclosion values (Exp.) for male and female flies of each genotype are noted in parentheses above the respective bars. b-d show photomicrographs of eyes of *ey-GAL4>Mof-RNAi* **(b)**, *ey-GAL4>Mof-RNAi/ EP3037* **(c)** and *ey-GAL4>Mof-RNAi/hsroj-RNAi* **(d)** male flies.

The few males eclosing following co-expression of *EP3037* allele of *hsrω* with *Mof-RNAi* transgene were sterile. Their testes and accessory glands appeared abnormal with accessory glands and seminal vesicle being smaller while sperms in testes were immotile (not shown).

### Reduction in eye size following *ey-GAL4* driven expression of *Mof-RNAi* transgene was partially rescued by over-expression of hsrω

Down-regulation of Mof in developing eye discs with the *ey-GAL4* driver resulted in smaller eyes (Fig. 4b) in all the male individuals but not in females (not shown). Following coexpression of *EP3037* allele of *hsrω*, the size of eyes was substantially restored in all the individuals (Fig. 4c). Interestingly, down-regulation of hsrω by expression of *hsrω-RNAi* in Mof depleted background did not restore the eye size (Fig. 4d)

### *Disruption of Megator distribution and DCC assembly following down-regulation of Mof transcripts were partially restored by over-expression of* hsrω

Previous studies have suggested Megator to be an interactor of DCC as well as *hsrω* (Zimowska and Paddy 2002; Vaquerizas *et al*. 2010, Singh and Lakhotia 2015). Therefore, the nuclear distribution of Megator and Msl-2 was examined in late male larval SG (Fig. 5a-h) and MT (Fig. 5i-p) of different genotypes. WT nuclei showed the Megator to be present, as reported earlier (Zimowska and Paddy 2002; Singh and Lakhotia 2015), in a prominent nuclear peripheral rim and as speckles in the nucleoplasm (Fig. 5a, i), while the Msl-2 was, as noted above, localized in a restricted area of the nucleus (Fig. 5e, m). In *Act-GAL4>Mof-RNAi* expressing male SG (Fig. 5b) and MT (Fig. 5j) cells, the peripheral rim of Megator was largely disrupted and the overall levels of nucleoplasmic Megator were also substantially reduced. Interestingly, about 75% of the examined nuclei (N=50) showed Msl-2 protein to be nearly absent in *Mof-RNAi* transgene expressing SG and MT nuclei while the rest showed a small presence of localized Msl-2 (not shown). Co-expression of *EP3037* allele of *hsrω* (Fig. 5c, g, k, o), but not of *hsrω-RNAi* transgene (Fig. 5d, h, l, p), with *Mof-RNAi* resulted in a partial but significant restoration of normal distribution of Megator as well as Msl-2 proteins.

**Fig. 5.**
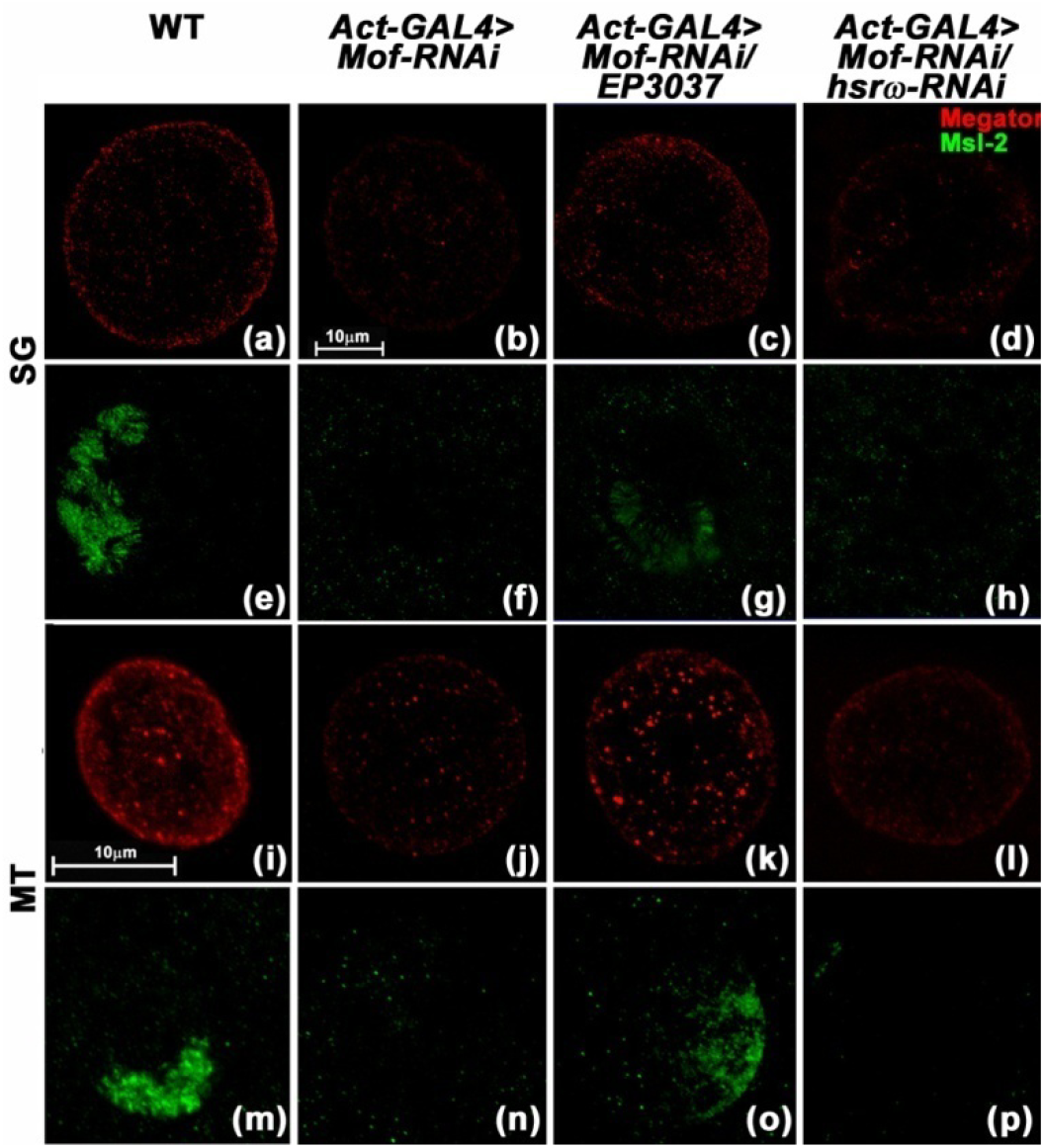
Expression of *Mof-RNAi* transgene affects distribution of Megator and Msl-2 in male nuclei which are partially restored by co-expression of *EP3037*. Confocal projection images show localization of Megator (red, a-d and i-l) and Msl-2 (green, e-h and m-p) in SGs (a-h) and principle cell nuclei of MTs (i-p) from late larvae of different genotypes, indicated on the top each column. Scale bar in (b) applies to images in a-h, while that in (i) applies to i-p.

## Discussion

Our present genetic study reveals that the long non-coding RNA gene *hsrω* interacts with the *Gcn5* and *Mof* genes since lethality and other phenotypes following down-regulation of Gcn5 or Mof were found to be partially suppressed by over-expression of hsrω transcripts. It is interesting that the male specific phenotypes associated with Mof, which is essential for hyperactivity of the X-chromosome in males (Hilfiker *et al*. 1997; Kind *et al*. 2008), were also partially suppressed by over-expression of hsrω. As noted in Introduction, both these HATs are involved in different aspects of chromatin remodeling and gene activity, with Mof, as a member of the DCC, being primarily responsible for H4K16 acetylation on male X chromosome to establish its hyperactive state (Conrad and Akhtar 2012). An earlier study (Onorati *et al*. 2011) noted interaction between Iswi, another chromatin remodeling component, and hsrω transcripts. However, unlike the partial suppression of Iswi-null phenotypes by down-regulation of hsrω transcripts observed by Onorati *et al*. (2011), over-expression of *hsrω* gene was found to partially suppress the Gcn5 and Mof phenotypes. These opposing effects of the *hsrω* gene activity seem to be related to the known antagonistic interactions between Iswi and Mof.

Corona *et al*. (2002) showed that following ectopic expression of Mof in Iswi depleted background in females, the paired X chromosomes also become bloated whereas in *mle* mutant background, Iswi depletion had no effect on male X-chromosome organization. Corona *et al*. (2002) also showed that co-depletion of Iswi and Mof exaggerated *Iswi* or *Mof* mutant phenotypes. Further, the DNA-bound basic patch of H4 is specifically bound by Iswi SANT domain but the state of Mof mediated acetylation of adjacent K16 lysine residues can negatively influence the Iswi remodeling functions by inhibiting its binding on chromosomes (Clapier *et al*. 2001, 2002; Corona *et al*. 2002; Hamiche *et al*. 2001). In view of these findings, it has been suggested (Corona *et al*. 2002) that the DCC-based H4K16 acetylation by Mof works antagonistically to the ATPase dependent compaction of chromatin by Iswi. Thus Mof down-regulation may permit greater association of Iswi on male X chromosome which would prevent its open chromatin conformation. Indeed the single X-chromsome in *Mof-RNAi* expressing male larval salivary glands appeared narrower and as darkly stained as the autosomes (not shown) instead of showing its characteristic enlarged width and pale staining. Since Iswi and hsrω transcripts interact physically and Iswi is required for the biogenesis of omega speckles (Onorati *et al*. 2011; Singh and Lakhotia 2015), it is likely that over-expression of *hsrω* may engage more of Iswi for omega speckle organization (Onorati *et al*. 2011; Singh and Lakhotia 2015), which may thus reduce its action on the male X-chromosome and consequently partially compensate for the reduced levels of Mof.

Megator, a nuclear matrix and nuclear pore complex protein, has been shown to have a role in the assembly of DCC (Mendjan *et al*. 2006; Akhtar and Gasser 2007; Vaquerizas *et al*. 2010). Megator also co-localizes with omega speckles (Singh and Lakhotia 2015) and binds with 93D locus following heat shock (Zimowska and Paddy 2002). A role of Megator in DCC assembly is also supported by our finding that the Megator levels and its localization get affected in Mof-down regulated cells. In view of the partial restoration of the levels and spatial location of Megator in cells that co-expressed *Mof-RNAi* and the *EP3037* over-expressing allele of *hsrω*, we speculate that the enhanced levels of hsrω transcripts may increase the stability and/or its availability for a more efficient assembly of the DCC on male X chromosome when Mof is limited.

Gcn5 acetylates histones at various positions and plays crucial role in *Drosophila* development (Kuo and Allis 1998; Carre *et al*. 2005). Gcn5 is important for Iswi activity since lysine 753 in the HAND domain of Iswi present in the NURF chromatin remodeling complex is required to be acetylated by Gcn5 for its activity (Ferreira *et al*. 2007). Thus down-regulation of Gcn5 would reduce acetylation of Iswi and thereby reduce its activity. Compromising the Iswi activity would affect, besides, many other things, the organization of the male X chromosome and the omega speckle biogenesis (Onorati *et al*. 2011; Singh and Lakhotia 2015). One of the paths that can partially compensate for reduced Iswi activity in the absence of Gcn5-mediated acetylation could be the improved stability/availability of Megator and DCC following over-expression of *hsrω*. Additionally, the substantial restoration of the levels of Gcn5 transcripts following overexpression of *hsrω* may mean that the Gcn5 functionality may not actually be much affected when *Gcn5-RNAi* transgene is co-expressed with the over-expressing *EP* allele of *hsrω*. The mechanism of up-regulation/stabilization of Gcn5 transcripts by over-expression of *hsrω* remains to be known.

The hsr transcripts are known to interact with diverse proteins involved in a variety of regulatory pathways including RNA processing, apoptosis, proteasomal activities, Ras signaling, Hsp83, nuclear matrix organization, chromatin remodellers etc, (Lakhotia 2011, 2016). Earlier studies from our laboratory (Mallik and Lakhotia 2010) had shown the hsrω transcripts to interact with CBP HAT. Present study adds two more HATs to the list of regulatory proteins that interact with these lncRNAs. It remains to be seen if these interactions are direct or indirect. Our other unpublished results indicate that the hsrω transcripts also interact with other members of the DCC like the Msl-1and Msl-2. Interactions with such diverse array of regulatory proteins confirm that some of the lncRNAs act as hubs in cellular regulatory networks (Arya *et al*. 2007; Lakhotia, 2011, 2012, 2015, 2016). The *hsrω* gene is now reported to generate seven lncRNAs, ranging in size from 1.2kb to ~21kb, through alternate transcription start and termination sites and regulated splicing of its single intron (Lakhotia 2016; www.flybase.org). Other studies in our laboratory (R. R. Sahu and S. C. Lakhotia unpublished) indicate that the 27aa encoding ORF in the 1.2kb hsrω transcript RC is translated. In view of the production of such diverse transcripts and a small polypeptide, it is not surprising that the *hsrω* gene impinges upon so many of the regulatory networks. The diversity of this gene’s products and their actions pose a big challenge to unravel modes and nature of interactions. However, it is expected that further studies with increasingly powerful approaches would gradually unfold the long and multi-pronged reach of lncRNAs.

## Acknowledgement

I am thankful to the Bloomington Stock Center for *Gcn5-RNAi* and *Mof-RNAi* fly stocks. I am grateful to Dr H Saumweber, Germany, and Dr. M Kuroda, USA, for generously providing the BX34 and anti-Msl-2 antibodies respectively. This work was supported by the Junior Research Fellowship and Senior Research Fellowship from the Council of Scientific and Industrial Research, India, and Senior Research Fellowship from Department of Biotechnology, India to DPC. The confocal microscope facility is funded by the Department of Science and Technology, India.

